# Cellular Senescence Impairs Tendon Extracellular Matrix Remodeling in Response to Mechanical Unloading

**DOI:** 10.1101/2023.12.22.572594

**Authors:** Emma J. Stowe, Madelyn R. Keller, Brianne K. Connizzo

**Affiliations:** Department of Biomedical Engineering, Boston University, Boston, MA 02215, United States

**Author notes:** Correspondence: Brianne K. Connizzo, Department of Biomedical Engineering, 44 Cummington Mall, ERB 403, Boston, MA 02215.

**Keywords:** Senescence, Aging, Extracellular Matrix, Tendon, Explant, Remodeling

## Abstract

Musculoskeletal injuries, including tendinopathies, present a significant clinical burden for aging populations. While the biological drivers of age-related declines in tendon function are poorly understood, it is well accepted that dysregulation of extracellular matrix (ECM) remodeling plays a role in chronic tendon degeneration. Senescent cells, which have been associated with multiple degenerative pathologies in musculoskeletal tissues, secrete a highly pro-inflammatory senescence-associated secretory phenotype (SASP) that has potential to promote ECM breakdown. However, the role of senescent cells in the dysregulation of tendon ECM homeostasis is largely unknown. To assess this directly, we developed an *in vitro* model of induced cellular senescence in murine tendon explants. This novel technique enables us to study the isolated interactions of senescent cells and their native ECM without interference from age-related systemic changes. We document multiple biomarkers of cellular senescence in induced tendon explants including cell cycle arrest, apoptosis resistance, and SASP production. We then utilize this *in vitro* senescence model to compare the ECM remodeling response of young, naturally aged, and senescent tendons to an altered mechanical stimulus. We found that both senescence and aging independently led to alterations in ECM-related gene expression, reductions in protein synthesis, and tissue compositional changes. Furthermore, MMP activity was sustained, thus shifting the remodeling balance of aged and senescent tissues towards degradation over production. Together, this demonstrates that cellular senescence plays a role in the altered mechano-response of aged tendons and likely contributes to poor clinical outcomes in aging populations.

## 1 Introduction

Musculoskeletal injuries present a significant clinical burden for aging patients, accounting for 25-30% total years lived with disability in elderly populations (Briggs et al., 2016). These injuries cause pain, decreased mobility, loss of independence, and reduced quality of life. Tendons are an important fibroelastic musculoskeletal tissue that connects muscle to bone, transmitting mechanical forces and storing energy to facilitate movement. Tendon injuries are especially common in aging populations. Rotator cuff tears, one of the most common conditions, were found to affect more than 50% of people over 80 years of age (Teunis et al., 2014). In addition to the high prevalence of tendon injuries, aged patients often demonstrate poor healing outcomes characterized by high rates of re-injury (Ackerman et al., 2017; Mienaltowski et al., 2016). It has been documented that both acute tendon injuries and tendinopathies are associated with a chronic degenerative state. However, the progression and biological drivers of age-related tendon degeneration are poorly understood.

It is widely accepted that dysregulation of balanced extracellular matrix (ECM) turnover plays a role in tendon degeneration. Primary tendon cells, tenocytes, are responsible for remodeling the ECM in response to changing mechanical loads, allowing the tissue to adapt and repair matrix damage. This process is a delicate cell-mediated balance of matrix clearance and synthesis that is essential in maintaining healthy tissue function. Tenocytes produce matrix-degrading enzymes, such as matrix metalloproteinases (MMPs), to clear damaged matrix. In parallel, cells synthesize new matrix proteins that are incorporated into the matrix and re-organized into functional tissue structure. Both exercise and disuse have been reported to initiate biological adaptations, with sustained exercise reportedly increasing tendon mechanical strength and matrix synthesis (Rooney et al., 2017) and disuse resulting in a decline in mechanical properties (Almeida-Silveira et al., 2000; Couppé et al., 2012). These cell-driven adaptations allow the tissue to meet changing mechanical demands. However, biological factors that alter cellular processes, such as aging or disease, can lead to dysregulation of ECM remodeling and put tissues at increased risk for injury.

While there are several well-established cellular changes associated with aging (López-Otín et al., 2023), one of the most promising candidates implicated in disrupting ECM remodeling is cellular senescence. Cellular senescence is a permanent state of cell arrest in response to external damage stimuli or stressors. Senescent cells are documented to be resistant to apoptosis and accumulate in many aged tissues, including tendon, likely due to impaired immune clearance with aging (Hawthorne et al., 2023; Kohler et al., 2013; Tsai et al., 2011; Zhang et al., 2023). Importantly, the senescence-associated secretory phenotype (SASP), which has been reported to include pro-inflammatory cytokines and MMPs, has the potential to degrade tissue matrix and induce neighboring cells to a senescent phenotype via secondary senescence (Coppé et al., 2010). While senescence is essential in development, protection against cancers, and wound repair, senescent cells are documented to have a negative effect during aging, leading to a decline of regenerative capacity, increased inflammation, and loss of function of aged tissues. More recently, senescent cells have been implicated in degenerative musculoskeletal diseases, such as osteoarthritis (Coryell et al., 2021).

Much of what we know about senescent cells and their biomarkers comes from isolated cell culture, including the first descriptions of senescence by Hayflick almost 60 years ago (Hayflick, 1965). *In vitro* induction of premature cellular senescence has been informative in identifying and characterizing the phenotype of senescent cells, as well as screening potential senotherapies that selectively target senescent cell populations (Chaib et al., 2022). Senescence can be induced *in vitro* with various stress or damage stimuli including extensive replication, oxidative stress, oncogene activation, mitochondrial dysfunction, and DNA-damage (González-Gualda et al., 2021). Commonly used DNA-damaging agents to induce senescence *in vitro* include radiation and chemotherapy drugs, such as doxorubicin (Copp et al., 2021; Kirsch et al., 2022). Senescence has been induced previously using bleomycin in rat patellar tendon stem cells (Nie et al., 2021), irradiation in murine fibroblasts (Saito et al., 2020), and dexamethasone in primary human tenocytes (Poulsen et al., 2014). However, as 2D cell culture alone does not replicate native cell-cell and cell-matrix connections necessary for studying ECM remodeling, the link between cellular senescence and dysregulation of ECM maintenance has not yet been fully elucidated.

Our lab utilizes a unique model of live tendon explants to directly assess ECM remodeling in real-time within an isolated tissue that maintains intact cell-ECM interactions. This novel technique enables the study of local cell biology in a more physiologically relevant context without interference from age-related systemic changes, such as immune responses and metabolic disease. Previous work by our group investigated age-related changes of murine flexor digitorum longus (FDL) tendon explants in response to an altered mechanical stimulus (Connizzo et al., 2020). Despite no age-related changes at baseline, we found that aged tendons exhibited reduced metabolism, proliferation, and matrix synthesis in response to mechanical unloading, indicating an altered mechano-response with aging. Accompanying this, we found an increase in senescence-associated gene expression (p16, p19, p53) in aged tendons, suggesting a previously unknown role of senescent cell populations on matrix remodeling. While additional studies by our lab and others have suggested a role for senescent cells in age-related ECM pathology, the ability of senescent cells to perform functional matrix remodeling is still largely unknown.

Therefore, the overarching goal of this study was to directly investigate the role of senescent cells in the dysregulation of tendon ECM homeostasis. Our first aim was to develop a model of induced cellular senescence in both primary murine tenocytes and flexor tendon explants to characterize the senescent phenotype of tendon cells and study the unique signature of senescent tenocytes within their native matrix environment. Utilizing our newly developed *in vitro* senescence explant model, our secondary objective was to compare the matrix remodeling response of young, naturally aged, and senescent-induced tendons to an altered mechanical stimulus. We hypothesized that both aged and senescent tendons would exhibit altered ECM remodeling following a mechanical unloading injury, with a shift to processes that promote ECM degradation over synthesis.

## 2 Methods

### 2.1 Sample Preparation

Primary tenocytes and tendon explants were obtained from FDL tendons harvested from 4-month old (young) and 20-24-month old (aged) male C57BL/6J mice. Aged mice were obtained from the NIH/NIA Aging Rodent Colony, and young mice were obtained from Jackson Laboratories. Primary FDL murine tenocytes were isolated as described previously (Paredes et al., 2020). Briefly, tendons were dissected, quickly dipped in 70% ethanol to kill surface cells, minced, and digested with 2mg/mL of collagenase type I (Gibco) and 1mg/mL of collagenase type IV (Worthington Biochemical) in low-glucose DMEM for 2 hours at 37°C on a rocking shaker. Ten FDLs were pooled for each biological replicate. Cell suspensions were filtered with a 70-μm cell strainer, plated at P0 in a T-25 culture flask at 1,200 cells/cm^2^ and expanded in culture until P1 when senescence was induced. FDL tendon explants were harvested as described previously (Connizzo et al., 2020). After tissue extraction, explants were washed in 1x PBS supplemented with antibiotics and placed in culture medium. For both cells and explants, culture medium consisted of low glucose Dulbecco’s Modified Eagle Media (DMEM) (1 g/L (Fisher Scientific)) with 10% fetal bovine serum (Cytiva), 100 units/ml penicillin G and 100 μg/ml streptomycin (Fisher Scientific). Culture medium was replaced every two days of culture. Tendon explants were cultured in stress deprived conditions with no mechanical stimulus for up to 14 days (Figure 1).

**Figure 1:**
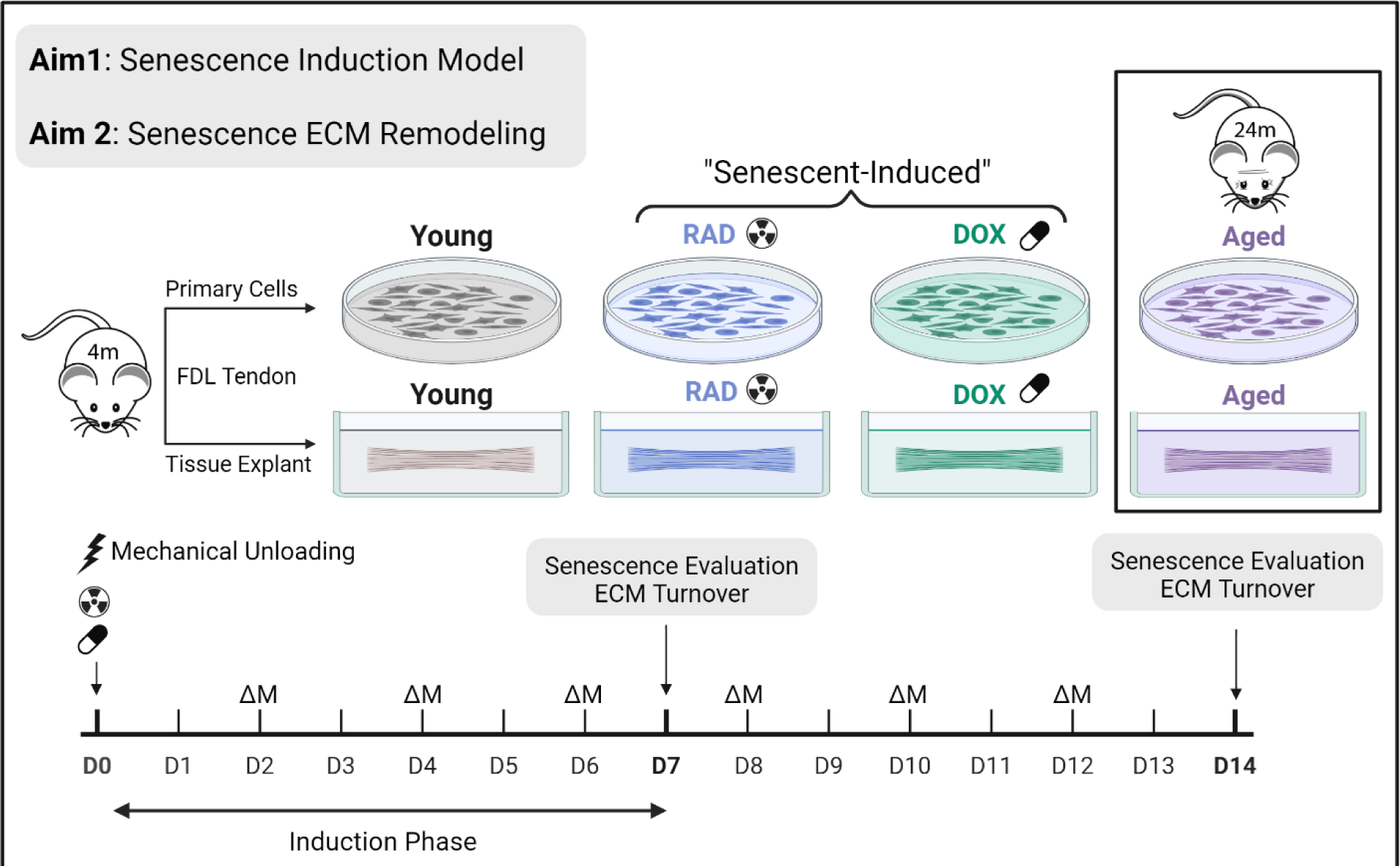
Experimental design. Primary cells and tendon explants were harvested from young mice. Senescence was induced with DOX and RAD treatment at D0. Aged tendons and cells were harvested from aged mice. Senescence induction and ECM turnover were evaluated after 7 and 14 days of culture under mechanical unloading. Media changes, ΔM, were performed every 2-days.

### 2.2 Senescence Induction

*In vitro* cellular senescence was induced in tendon explants and primary tenocytes with two methods: irradiation (RAD) and doxorubicin (DOX) (Figure 1). For RAD treatment, a 10Gy dose of irradiation was delivered at Day 0 of culture using an x-ray irradiator (Precision X-Ray). For DOX treatment, explants and cells were cultured in 200nM doxorubicin hydrochloride (Sigma) in culture medium for the first 72 hours, after which they were switched to standard medium for the remaining culture period. Senescence was evaluated after 7 days, and additional assays were performed at day 14 to confirm a sustained phenotype. It is well established that there is no universal biomarker for cellular senescence. Therefore, we evaluated senescence with a panel of biomarkers including DNA damage (γH2AX, Section 2.6), cell cycle arrest (thymidine incorporation, Section 2.4 and gene expression of cell cycle regulators, Section 2.5), secretory phenotype (protein secretion in media, Section 2.8), apoptosis resistance (Section 2.7), metabolic changes (Section 2.3), increased lysosomal content (SA-β-Gal expression, Section 2.6), and nuclear changes (loss of nuclear protein lamin B1, Section 2.5).

### 2.3 Explant Viability and Metabolic Activity

Explant viability was assessed using live/dead imaging (n=4/group). Whole explants were incubated for 5 minutes in 1x PBS with fluorescein diacetate (FDA; 4mg/ml, Sigma) and propidium iodide (PI; 1mg/ml, Sigma), to label viable and non-viable cells, respectively. Samples were mounted between two pieces of #1.5 coverslip with a small layer of 1xPBS and imaged via confocal microscopy (10x, Olympus FV3000). Multi-Area Time Lapse (MATL) software was used to scan across the desired region of interest to form a full tendon viability map. Z-stacks of approximately 80-100μm thickness were generated and maximum intensity projections (MIPs) were formed. The total number of live and dead cells was quantified with a custom MATLAB script that thresholds channels individually and analyzes the total number of cells in a 7mm region of interest (ROI). Viability is expressed as a percentage of number of live cells divided by total number of cells in the specified ROI. Cell density is calculated as the total number of cells divided by the ROI area.

Explant and cell metabolism were assessed throughout culture with a resazurin reduction assay (Rodríguez-Corrales & Josan, 2017). Samples (n=5/group/time) were incubated with media containing 10% v/v resazurin. After 3 hours, fluorescence intensity in spent medium was measured at 554/584 nm. Data is expressed normalized to control medium with resazurin, such that a value of ‘1’ represents no detectable metabolic activity.

### 2.4 Cell Proliferation, Matrix Biosynthesis, Matrix Composition

Synthesis rates of sulfated glycosaminoglycans (sGAG), total protein (indicative of collagen synthesis), and DNA (cell proliferation) were measured via incorporation of radiolabeled sulfate, proline, and thymidine, respectively. Radiolabels ^35^S-sulfate (2 μCi/mL), ^3^H-proline (1 μCi/mL), and ^3^H-thymidine (1 μCi/mL) (Perkin-Elmer) were added directly to culture medium for 24-hours (proline and thymidine radiolabeling performed in separate experiments). Following radiolabel incorporation, tendon wet weights were measured after soaking the tendon in 1xPBS for 1-minute. Tendons were then lyophilized for 3-hours and tendon dry weights were recorded. Water content of the tendon was calculated as the difference in wet weight and dry weight divided by the dry weight and multiplied by 100. Tendons (n = 5/group/time) were then digested with proteinase-K (5 mg/ml) (Sigma) for 18 hours and stored at -20C until further assays could be performed. Cell suspensions (n = 5/group) were lysed with sodium hydroxide and stored at -20C. Radiolabel incorporation was measured with a liquid scintillation counter (Perkin-Elmer) and values represent the rate of incorporation over the 24-hour time period. Incorporation rates for each explant sample are normalized to the tissue dry weight to account for variability in sample size. Incorporation rates for cells are normalized to the total number of cells.

Additional biochemical assays were then performed with the digest to quantify explant composition (Connizzo et al., 2020). sGAG content was measured using the dimethyl methylene blue (DMMB) assay. Double-stranded DNA content was measured using the PicoGreen dye binding assay. A 100 µL portion of each digest was then hydrolyzed using 12 M HCl, dried, resuspended, and assayed to measure total collagen content using the hydroxyproline (OHP) assay.

### 2.5 Quantitative Gene Expression

Explants from each group were collected at day 0 (baseline), day 7, and day 14 (n = 5/group/time). Explants were immediately flash-frozen with liquid nitrogen and stored at −80°C until RNA extraction, performed as described previously (Grinstein et al., 2018). Total RNA was then reverse-transcribed to cDNA and real-time polymerase chain reaction (PCR) was performed using the Applied Biosystems StepOne Plus RT-PCR with SYBR Green Master Mix (Applied Biosystems). Murine gene names and primer sequences are listed in the Supporting Information (Table S1). Briefly, we measured markers of cell cycle (p16^ink4a^, p53, p21), nuclear membrane (lamin B1), apoptosis (caspase-3), inflammation (IL-6, CXCL1, IL-1β), matrix metalloproteinases (MMP-1, MMP-3, MMP-13), and ECM proteins (collagen 1, decorin, fibromodulin). Expression for each gene was calculated from the threshold cycle (*C*_t_) value and fold changes were calculated by normalizations to the housekeeping gene, beta-actin, and to day 0 values using the double delta CT method (Livak & Schmittgen, 2001). All data are presented in log space.

### 2.6 Histological Assessment

Samples were analyzed for SA-β-Gal, **γ**H2AX, and p21 protein using standard immunohistochemical techniques. Fresh tendons (n = 5/group) were removed from culture, embedded in OCT media, and flash frozen. Frozen samples were transferred to -20°C until sectioning. SA-β-Gal samples were stored for a maximum of 72-hours before sectioning and fixing to prevent loss of activity. Tendons were cryosectioned with a 10μm thickness at -20°C. Cryosections were allowed to briefly air dry before being washed with 1x PBS and fixed for 10-minutes at room temperature with 4% paraformaldehyde. Cells were plated on 12mm #1 glass coverslips pre-coated with poly-d-lysine (Neuvitro Corporation) at a seeding density of 75,000 cells per 24-well, allowed to adhere overnight, and fixed for 10 minutes at room temperature with 2% paraformaldehyde. Fixed cells and tissue sections were washed with 1% BSA to remove fixation solution.

SA-β-Gal was stained using CellEvent™ Senescence Green Detection Kit (Invitrogen), following manufacturer guidelines. Immunostaining for **γ**H2AX (phospho-H2AX (Ser139) Alexa Fluor 647 conjugate antibody, Cell Signaling, 1:2000 dilution) and p21 (rabbit monoclonal to p21(ab188224), 1:500 dilution; goat anti-rabbit IgG H&L (Alexa Fluor® 647) (ab150079), 1:1000 dilution) were performed using standard immunostaining protocols with a 10-min permeabilization step in 0.2% Triton X-100. Stained tissue sections and cell cover slips were mounted with DAPI antifade mounting media (ProLong™ Gold Antifade Mountant with DAPI, Life Technologies) and imaged on an inverted microscope (10-20x, Olympus IX83) with appropriate filters (DAPI/FITC/Cy5). Small z-stacks were collected for tissue sections and 3D deconvolution was performed to improve resolution and contrast. MIPs were analyzed using a custom MATLAB script that thresholds channels individually and counts the number of cells in the FITC and Cy5 channels that co-localize with DAPI signal. For cells, single slice images were obtained, and background was subtracted from each image. A custom MATLAB script was utilized to determine significant signal overlap based off global threshold values. In cells, individual **γ**H2AX foci could be resolved and the number of foci per nuclei was quantified by counting puncta within each thresholded DAPI area. Since SA-β-Gal and **γ**H2AX were co-stained, dual staining was quantified by co-localizing DAPI with both FITC and Cy5 signals.

### 2.7 Apoptosis Resistance

Apoptosis was induced with 1μM staurosporine (SP) treatment for 24-hours. Repeated resazurin reduction assays were performed before and after treatment such that cell death due to SP could be directly monitored for each sample (n = 5/group). Percent loss in metabolism between repeated assays was calculated as a measure of cell death, where reduced cell death (reduced loss in metabolism) represents a resistance to apoptosis.

### 2.8 Secretory Profile Analysis

Activity of MMPs (1, 2, 3, 7, 8, 9, 10, 13, 14) was determined via analysis of spent culture medium (n = 5/group/time) using a commercially available FRET-based generic MMP activity kit (SensoLyte 520 Generic MMP Activity Kit Fluorimetric; Anaspec). MMP activity is represented as the concentration of MMP cleaved product (5-FAM-Pro-Leu-OH), the final product of the MMP enzymatic reaction. Secretion of SASP-associated proteins was assessed in spent media samples (n=5/group/time) using a custom multiplex immunoassay (UPLEX Biomarker Group 1 (mouse); Meso Scale Discovery) for 10 analytes: GM-CSF, IFN-γ, IL-1β, IL-6, IL-10, IL-13, KC/GRO (CXCL1), MCP-1, MIP-1α, and TNF-α.

### 2.9 Statistical Evaluation

All data are presented as mean ± 95% confidence interval. Data points more than two standard deviations from the mean were removed as outliers. Statistical evaluation was performed using one-way analysis of variance (ANOVA) at each time-point. Bonferroni corrected post hoc *t*-tests were then used to identify differences within each time point where appropriate. For all comparisons, significance was noted at \**p* < 0.05 and a trend set at *p* < 0.10.

## 3 Results

### 3.1 Senescence Induction in Primary Tenocytes

Cellular senescence was induced in primary murine tenocytes using radiation and doxorubicin treatment (Figure 1). After 7 days of senescence induction, the induced cells were compared to populations of young and naturally aged tenocytes. Representative images of tenocytes co-stained with DAPI, SA-β-Gal, and **γ**H2AX are shown in Figure 2a. Image quantification revealed increased SA-β-Gal+ and **γ**H2AX+ cell populations in both the aged and induction groups (RAD and DOX) compared to the young cells (Figure 2b, c). Additionally, a significantly increased number of cells co-staining for both SA-β-Gal and **γ**H2AX was observed in the aged, RAD, and DOX treatment groups (Figure 2d). Furthermore, both induction groups had a larger number of **γ**H2AX foci per cell, indicating increased accumulation of irreparable DNA damage (Figure 2e). The number of **γ**H2AX foci per cell was not increased in aged cells compared to young control cells. Significant decreases in cell proliferation were found in aged and DOX cells when compared to control cells, and although not significant and highly variable, RAD cells also exhibited a reduction in proliferative capacity (Figure 2f). Metabolic activity also appeared to decrease with RAD and DOX treatment, although statistical significance could not be achieved with our sample size (Figure 2g).

**Figure 2:**
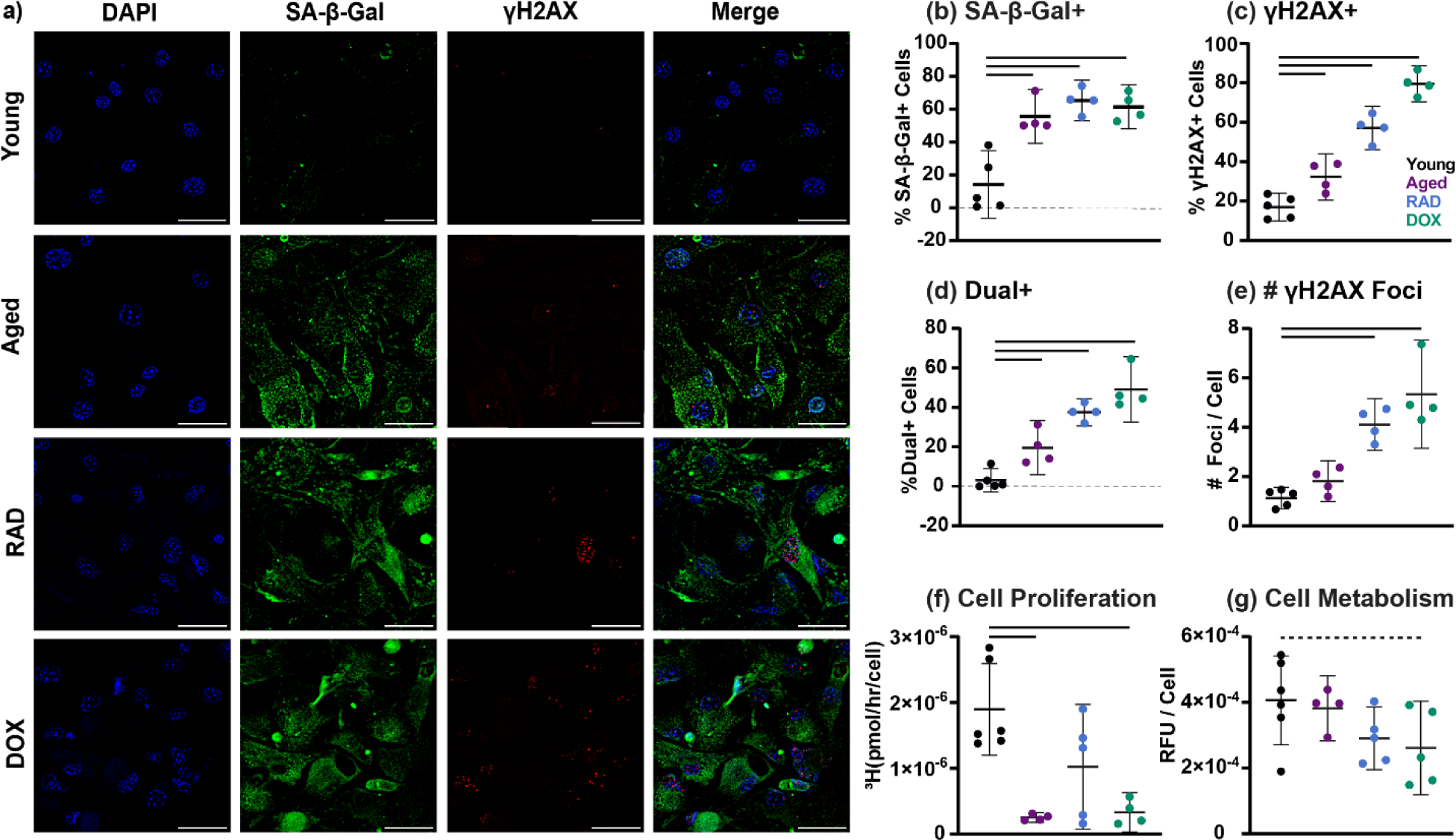
Senescence Induction in Murine Tenocytes. (a) Representative images of young, aged, RAD, and DOX cells stained with DAPI (blue), SA-β-Gal (green), and γH2AX (red). Associated image quantification for (a) SA-β-Gal+ cells (b) γH2AX+ cells (c) dual label+ cells and (d) number of γH2AX foci per cell. (e) Total cell proliferation and (f) metabolic activity. Scale bars are 50μm. Significance set at p<0.05 (solid lines) and trends at p<0.1 (dashed lines) compared to young cells.

### 3.2 Senescence Induction in Tendon Explants

Following validation of senescence-induction methods and senescence-assessment assays in primary murine tenocytes, senescence was induced in live tendon explants with both RAD and DOX treatment. First, we confirmed all groups were viable after 7 and 14 days of culture (Figure S1). We then assessed a time course of explant proliferation as a metric of cell cycle arrest. For a young explant under stress deprivation, proliferative capacity increased throughout culture until day 7 (Figure 3a). Compared to the young tissues, both DOX and RAD treated explants exhibited significantly reduced proliferation (Figure 3a). This phenotype started at day 5 and continued out to day 14. At day 7, RAD and DOX tendons also exhibited significant reductions in metabolic loss following staurosporine treatment, indicating resistance to apoptotic cell death (Figure 3b).

**Figure 3:**
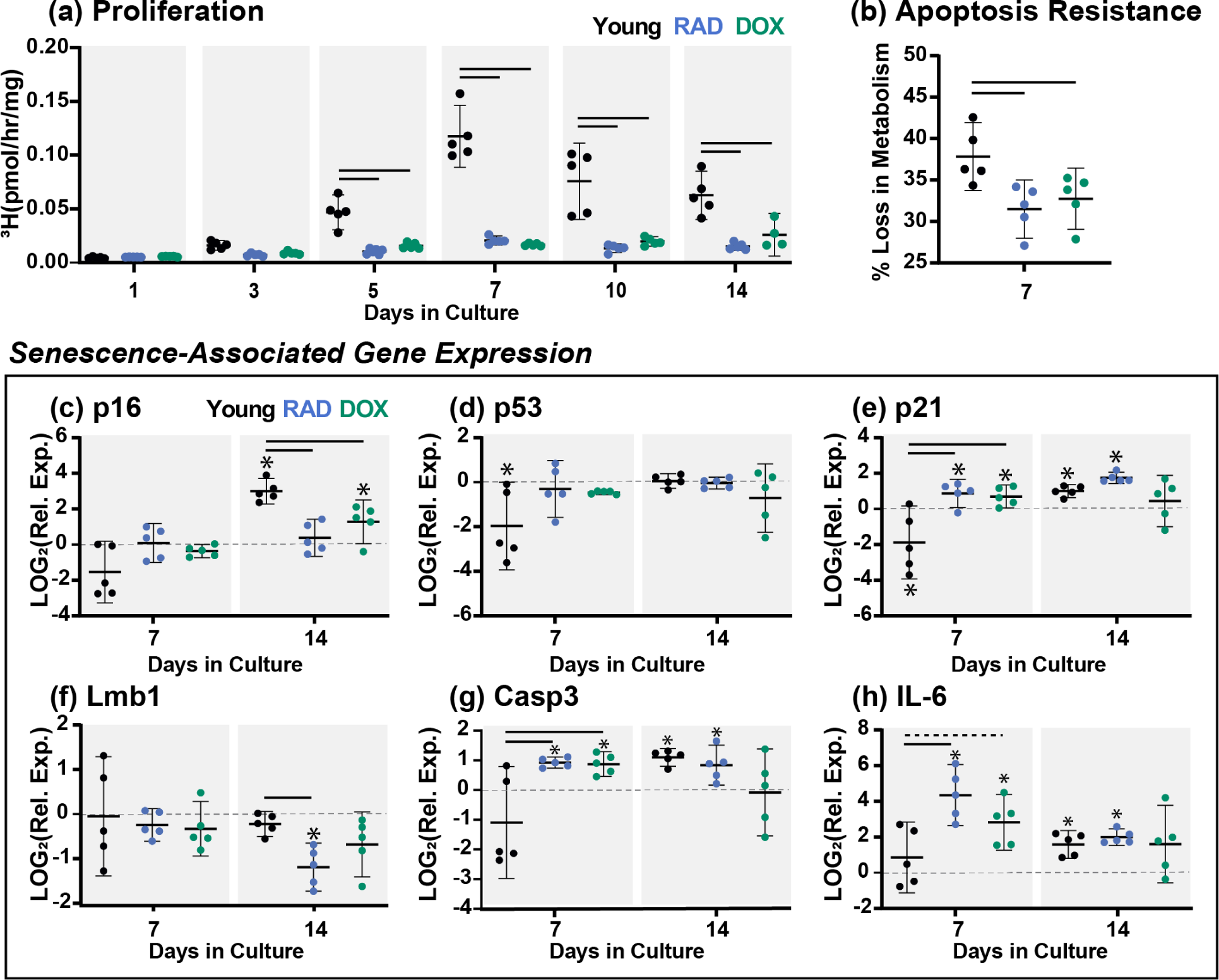
Senescence Induction in Tendon Explants. (a) Cell proliferation at days 1, 3, 5, 7, 10, and 14. (b) Apoptosis resistance at day 7 shown as percent loss in metabolism following 24-hr treatment with staurosporine. Senescence-associated gene expression at day 7 and day 14 for (c) p16, (d) p53, (e) p21, (f) Lmb1, (g) Casp3, and (h) IL-6. Gene expression data presented as relative expression to D0. Significance set at p<0.05 (solid lines) and trends at p<0.1 (dashed lines) compared to the young group. * indicates significant difference from day 0.

Senescence-associated gene expression was assessed at day 7 and day 14 (Figure 3c-h). No significant differences were seen in p16 expression at day 7 (Figure 3c). By day 14, young and DOX explants upregulated p16 compared to day 0. Consequently, RAD and DOX explants expressed significantly lower p16 than young explants. At day 7, p53 expression was downregulated from baseline in young explants but no different from day 0 in senescent-induced explants (Figure 3d). No differences in p53 were seen between groups at day 14. Interestingly, p21 was downregulated at day 7 in young explants but upregulated in the RAD and DOX groups, with significant differences between groups (Figure 3d). However, these differences were lost by day 14 as young explants also exhibited a significant upregulation of p21 relative to baseline. No differences in Lmb1 were seen at day 7 (Figure 3f). However, by day 14 RAD tendons exhibited a downregulation in Lmb1 compared to young tendons. At day 7, Casp3 gene expression was increased for RAD and DOX explants compared to the young group (Figure 3g). Between days 7 and 14, young explants undergo significant increases in Casp3 expression, resulting in no significant differences between groups at day 14. IL-6 was found to be upregulated for RAD and DOX explants, compared to the young group, at day 7 (Figure 3h). However, again, these differences were lost by day 14 when young explants exhibited significant upregulation of IL-6 relative to baseline.

Histological assessment for SA-β-Gal, p21, and **γ**H2AX was performed at day 7 only. Representative images for SA-β-Gal staining are shown in Figure 4a. Quantification revealed significant increases SA-β-Gal between D0 and D7 for all groups (Figure 4b). However, no significant differences were found at D7 between control and senescent-induced groups. Representative p21 immunostaining is shown in Figure 4c. Quantification of p21 staining showed increased p21 protein expression for RAD tendons at day 7 compared to both young and DOX groups (Figure 4d). Surprisingly, **γ**H2AX analysis showed little staining (<10% positive cells) for all groups at both D0 and D7 (Figure S2).

**Figure 4:**
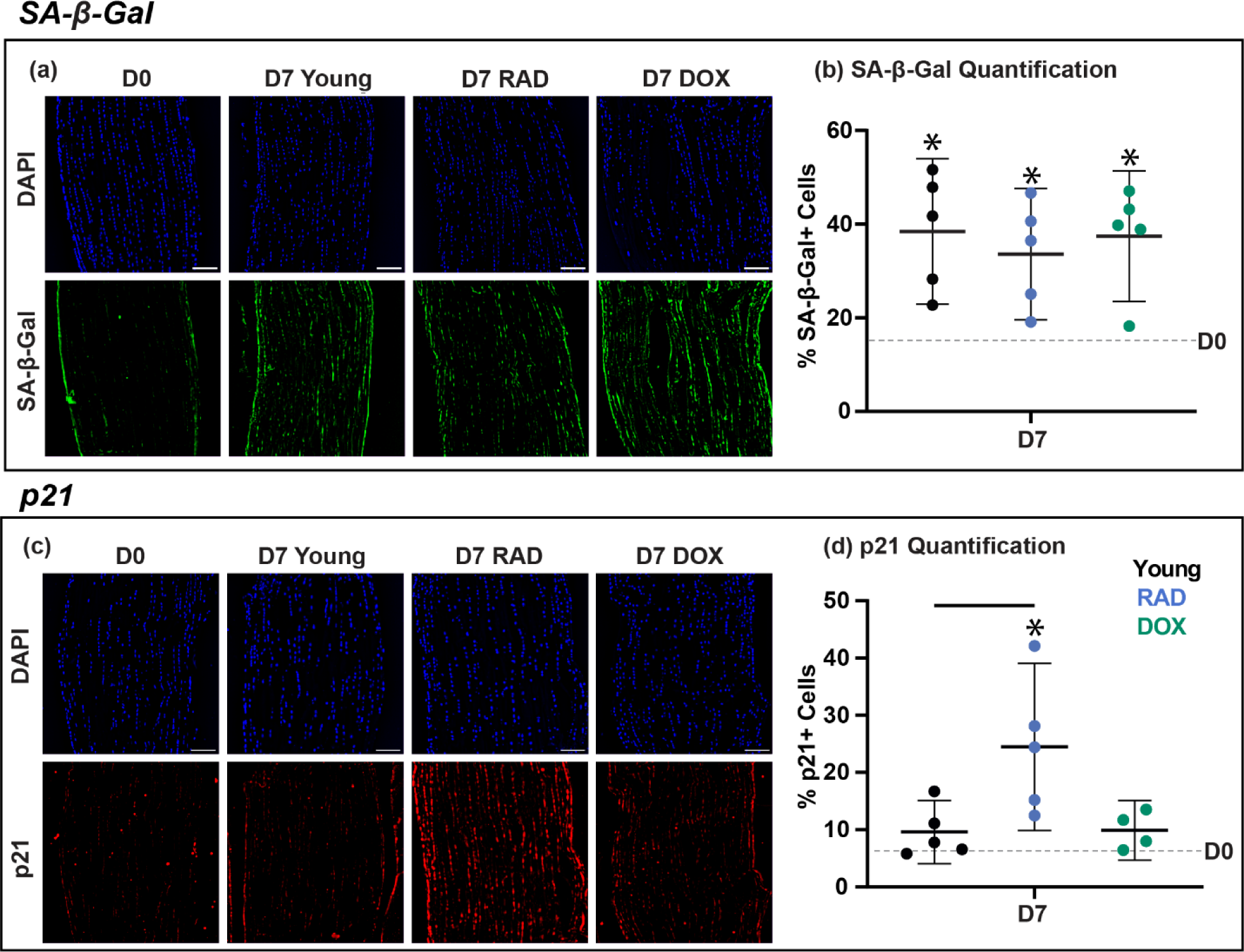
Senescence Induction in Tendon Explants. (a) Representative images and associated quantification (b) of young, RAD, and DOX tendon explants stained with DAPI (blue) and SA-β-Gal (green) at day 7. (c) Representative images and associated quantification (d) of young, RAD, and DOX tendon explants stained with DAPI (blue) and p21 (red) at day 7. Scale bars are 50μm. Significance set at p<0.05 (solid lines) compared to young group. * indicates significant difference from day 0.

Senescence in aged tendons was assessed via comparison of baseline metrics in freshly harvested young and aged tissues. Significant increases in gene expression of p16, p53, p21, IL-6, MMP-1, and MMP-3 were found in aged tissues compared to young tissues (Figure S3). However, no differences were found in the number of cells staining positively for SA-β-Gal, p21, and **γ**H2AX (Figure S2 and Figure S3).

### 3.3 Cellular Senescence and ECM Turnover

With our senescence-induction model validated, we next aimed to assess the capacity for ECM remodeling following an altered mechanical stimulus in young, aged, and senescent tendon explants. At baseline, aged tendons expressed reduced collagen 1, fibromodulin, and decorin compared to young tendons (Supplemental Figure 3). At day 7, collagen 1 expression was significantly downregulated from baseline for RAD and DOX explants, but it was no different from day 0 for young and aged tendons (Figure 5a). By day 14, collagen 1 expression was upregulated from baseline in all groups and induced senescent groups exhibited significant decreases in expression compared to the young group. At day 7, while decorin expression was found to be downregulated from baseline in all groups, aged, RAD, and DOX explants exhibited significant increases in expression compared to the young group (Figure 5b). Between days 7 and 14, young tendons had increased decorin expression. By day 14, induced senescent groups had decreased decorin expression compared to the young group. While all groups had significant downregulation of fibromodulin at day 7, aged and RAD groups exhibited higher expression than young tendons (Figure 5c). By day 14, fibromodulin expression was now found to be no different than day 0 in young tendons. However, aged and senescent tissues had reduced fibromodulin expression compared to young tissues.

**Figure 5:**
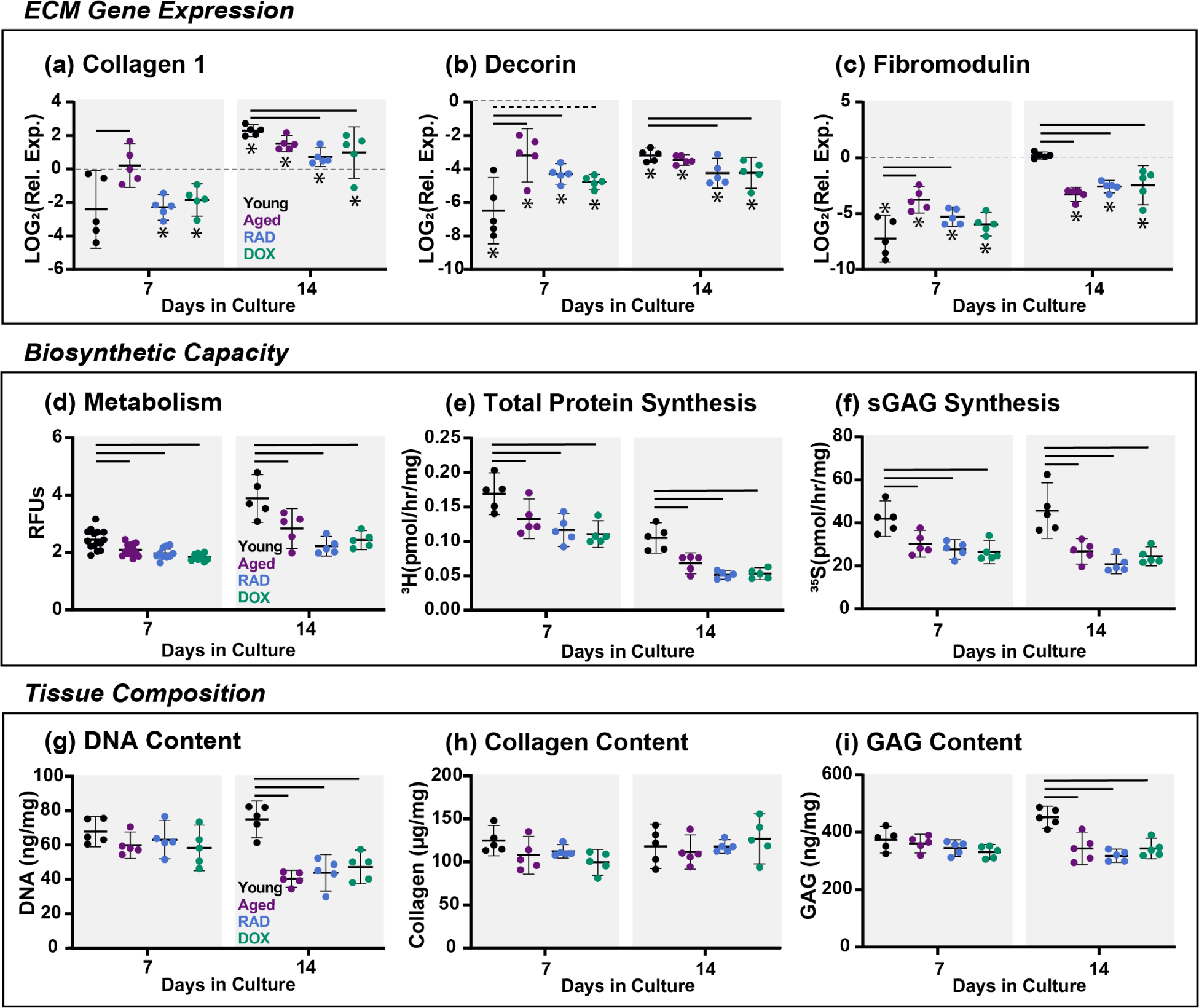
Matrix Synthesis and Composition. Relative gene expression of (a) collagen 1, (b) decorin, and (c) fibromodulin. Biosynthetic capacity assessed via (d) explant metabolic activity and ECM synthesis rates for (e) total protein and (f) sGAGs. Tissue composition of (g) DNA, (h) collagen, and (i) GAG. All data at day 7 and day 14. Significance set at p<0.05 (solid lines) and trends at p<0.1 (dashed lines) compared to the young group. * indicates significant difference from day 0.

We next examined the biosynthetic capacity of aged and senescent tendon explants. At both day 7 and day 14, explant metabolism was reduced in aged, RAD, and DOX groups compared to the young explants (Figure 5d). Furthermore, aged and induced senescent explants had significantly lower total protein synthesis compared to young tendons at both day 7 and day 14 (Figure 5e). Similarly, sGAG synthesis was decreased for senescent and aged tendons at day 7 and day 14 compared to young tendons (Figure 5f). Finally, we assessed tissue composition for young, aged, and senescent tendons after 7 and 14 days. No changes in tissue composition were found at day 7 (Figure 5g-i). However, by day 14, there were significant decreases in DNA content in aged, RAD, and DOX explants compared to the young group (Figure 5g). Additionally, aged and senescent tendons had significant reductions in total GAG (Figure 5i) and water content (data not shown) at day 14.

To get a holistic picture of tissue remodeling, we also assessed ECM breakdown via the production of MMPs and secreted inflammatory proteins for young, aged, and senescent tissues. On the gene expression level, MMP-1 expression was decreased in aged and senescent tissues compared to young tissues, with differences that become more pronounced by day 14 (Figure 6a). Conversely, MMP-3 and MMP-13 were both significantly upregulated from day 0 for all groups at both timepoints (Figure 6b, c). While aged tendons had reduced expression of MMP-3 compared to young, induced senescent explants had increased MMP-3 expression at day 7 (RAD & DOX) and 14 (DOX only) (Figure 6b). Both aged and DOX tendons had significantly higher MMP-13 expression than young tissues at day 7 (Figure 6c). By day 14, only aged tissues had increased MMP-13 expression compared to young. To get a full picture of time-dependent changes, we assessed MMP activity and inflammatory protein secretion at 2,7-, and 14-days after the initiation of the stress deprivation injury. For young tendons, generic MMP activity increased throughout the 14-day culture (Figure 6d). MMP activity in aged and senescent tissues was on the same levels as young tissues, with no significant differences between groups. Similarly, the production of IL-6 (Figure 6e), MIP-1α (Figure 6f), MCP-1 (Figure 6g), IL-13 (Figure 6h), and TNFα (Figure 6i) increased throughout culture for all groups, with little significant differences between groups. Conversely, the production of KC-GRO (Figure 6j) and GM-CSF (Figure 6k) decreased throughout culture. However, there were still no significant differences between groups.

**Figure 6:**
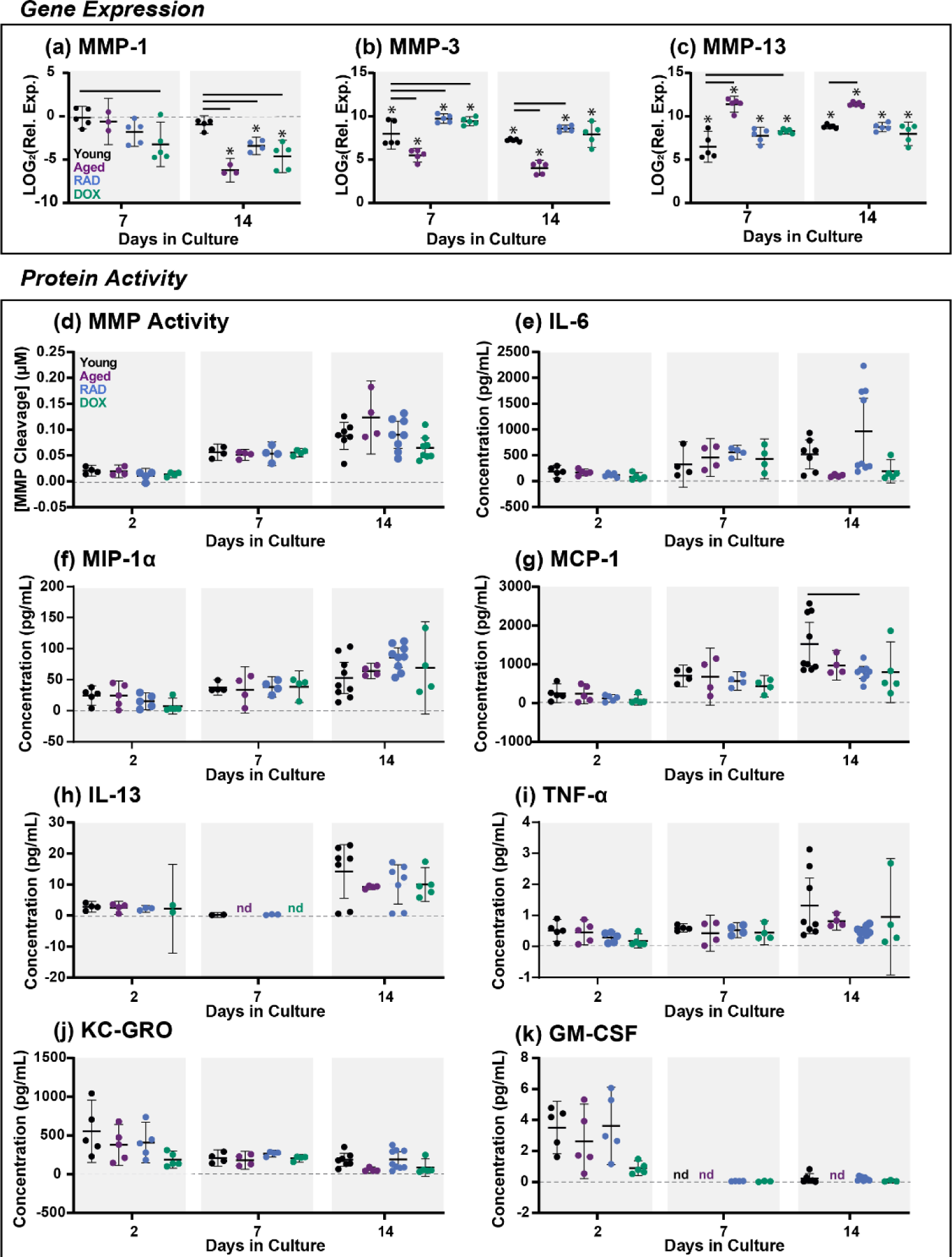
Matrix Degradation. Gene expression of (a) MMP-1, (b) MMP-3, and (c) MMP-13. (d) Generic MMP protein activity and concentration of secreted inflammatory proteins in culture media at days 2,7, and 14. (e) IL-6 (f) MIP-1α, (g) MCP-1 (h) IL-13 (h) TNF-α, (j) KC-GRO, and (k) GM-CSF. “nd” is listed where concentrations were below the detection limit of the assay and therefore not detected. Significance set at p<0.05 (solid lines) compared to young explants. * indicates significant difference from day 0.

## 4 Discussion

The objectives of this study were two-fold: (1) to establish a model of induced cellular senescence in murine tenocytes and tendon explants, and (2) to use this model to investigate how aging and cellular senescence independently affect the capacity for cell-driven matrix remodeling. We started by inducing cellular senescence in 2D cell culture, as this is more commonly done in the literature (Copp et al., 2021; Kirsch et al., 2022; Nie et al., 2021; Poulsen et al., 2014). It is well established in the field that there is no universal biomarker for cellular senescence and the senescent phenotype appears heterogeneous and dynamic across cell types and induction stimuli (Hernandez-Segura et al., 2018). Thus, it is of high importance to characterize the unique senescent signature of various cell types, including tenocytes. Senescence is typically evaluated with a panel of general hallmarks and biomarkers including DNA damage (ex. **γ**H2AX), cell cycle arrest, secretory phenotype production, apoptosis resistance, metabolic changes, morphological changes, SA-β-Gal expression, and nuclear changes (González-Gualda et al., 2021). It is important to note that while all these biomarkers have been observed in senescent cells, they are not necessarily present in all senescent cells. Due to this, a concurrent validation of multiple hallmarks is recommended to confirm a senescent phenotype *in vitro*.

Using this framework, we have documented successful induction of cellular senescence in primary murine tenocytes using two methods: 10Gy irradiation and 200nM doxorubicin treatment. We have shown that senescent tenocytes exhibit common biomarkers of cellular senescence including SA-β-Gal, **γ**H2AX, and cell cycle arrest, and this is consistent with induced senescent tendon cells across the field (Cai et al., 2022; Nie et al., 2021; Poulsen et al., 2014). Our approach resulted in an induction efficiency of 37% for RAD and 49% for DOX treated cells, based on the percentage of dual-positive cells. Reported induction efficiencies in the literature vary, with over 75% of rat patellar cells after 5-days of treatment with bleomycin (Nie et al., 2021) and around 30% of human tenocytes following both 3 and 7 days dexamethasone treatment (Poulsen et al., 2014). Importantly, we found low baseline levels of senescence (3%) in young murine tenocytes and elevated levels (20%) in aged murine tenocytes. In addition, the number of aged cells staining positive for SA-β-Gal is similar to what has been documented previously in the literature in tendon stem cells (M. Chen et al., 2020). Together, our work along with previous data demonstrates that aged tendon cell populations contain senescent cells and young tendon cells can be induced to senescence with various stress stimuli.

We next aimed to induce cellular senescence in tendon cells within their native ECM. Using the same induction methods (RAD and DOX) with a slightly larger panel of assessment biomarkers, we have documented successful induction of cellular senescence in live murine tendon explants. Most convincingly, we see substantial reductions in cellular proliferation, indicative of cell cycle arrest, beginning as early as day 5 and sustaining throughout the culture period. Additionally, we document resistance to apoptotic cell death, elevated expression of cell cycle regulators, reduced expression of nuclear proteins, and increased expression of SASP factors. Interestingly, expression of p21 was elevated in senescent groups at the gene and protein (RAD only) levels, suggesting p21 as a key marker of cellular senescence at this timepoint in our model system. On the protein level, senescent explants demonstrate sustained production of MMPs and other inflammatory SASP analytes, but on the same level as young explants. Similarly, SA-β-Gal expression, while increased from baseline levels, was no different between young and senescent groups. However, because we are simultaneously assessing compounding the effects of senescence induction and a mechanical unloading injury (stress-deprivation), the interpretation of individual biomarkers is more complex.

It is clear that the mechanical unloading injury alone has substantial impact on cell health at late stages of culture. Specifically, we observe declines in tissue viability and cell proliferation, accompanied by increased expression of injury and apoptosis markers, increased MMP activity, and secretion of inflammatory factors. In alignment with previous work (Egerbacher et al., 2008; Leigh et al., 2008), these biomarkers support a highly-inflammatory and injurious response to mechanical unloading causing apoptosis and cellular degradation in young tissues. Surprisingly, we also found significantly increased SA-β-Gal, p16, and p21 in young tissues compared to freshly harvested tissues. It is possible that the mechanical unloading could induce some cells to cellular senescence via an injury-initiated mechanism. Previous work has reported *in vivo* senescence induction (reduced cell division, p19, p53) in patellar tendons following injections of Botox to induce mechanical stress-deprivation (P. Chen et al., 2021). Many other groups are also interested in this idea of injury-initiated cellular senescence, particularly in the context of wound healing (Samdavid Thanapaul et al., 2022). However, there is still much debate about the interplay of senescent cells and tissue healing, particularly the permanent nature of these observed senescent phenotypes (Chu et al., 2020). Additionally, we did not find cell cycle arrest in concert with other senescence-associated changes leading to questions regarding the distinctions between senescence, quiescence, and early apoptosis. Stress-deprivation alone clearly has pronounced effects on the expression of multiple biomarkers of cell health, making it difficult to tease out the phenotype of senescence induction from that of chronic mechanical injury. Future studies will remove this confounding variable by using custom-designed tensile loading bioreactors to perform senescence induction under physiologic levels of mechanical loading.

By comparing the senescence induction efficiencies and phenotypes between primary cells and tendon explants, we can assess the role of the extracellular matrix in the induction of cellular senescence. While we did not assess the exact same markers in cells and tissue explants, we do see SA-β-Gal expression and reduced proliferation in both model systems. Our quantification showed 60% SA-β-Gal+ cells and high expression of **γ**H2AX (>60%) in induced primary tenocytes and less than 40% of SA-β-Gal+ cells and little **γ**H2AX (<15%) in induced tendon explants. This could suggest a potential protective mechanism of the ECM towards senescence induction that has not been explored due to the lack of tissue induction studies. However, it’s important to note that assessment is different in whole tissues and isolated cells due to the difficulties working with a dense, collagenous matrix. Therefore, in the future we plan to remove cells from the senescent induced tendon matrix and compare to those induced in 2D culture with the same assessment methods for a more accurate understanding of this potential phenomena. Furthermore, it appears DOX treatment may be more efficient in inducing senescence in primary cells while RAD treatment results in higher expression of senescence biomarkers in tendon explants. Again, this highlights the heterogeneous profile of senescent cells with specific phenotypes that depend on the model system, cell type, induction stimuli, and timepoint (González-Gualda et al., 2021).

After characterizing our senescent explant model, we then began to assess how cellular senescence and natural aging independently affect the capacity for ECM remodeling following a mechanical unloading injury. We first note some relevant findings regarding cell health and cellular activity that could affect matrix turnover. We documented reduced DNA content in senescent tissues at the end of the culture period. While this typically is attributed to reduced cell numbers, we did not find a corresponding change in cell viability. However, our DNA content assay is a quantification of double stranded DNA (Singer et al., 1997), and therefore it’s possible this reflects senescence-associated DNA damage in induced groups. We also found significant declines in metabolic activity of senescent tissues at both timepoints. The interactions of metabolic processes with both senescence and ECM remodeling are heavily studied areas of research (Sullivan et al., 2018; Wiley & Campisi, 2016); while these complex processes are largely outside of the scope of these current findings, we recognize altered metabolic activity is commonly observed with cellular senescence and could contribute to ECM remodeling outcomes.

Regardless, using a multi-scale assessment of matrix turnover we document that senescent tendons exhibit a decreased capacity to synthesize new ECM in response to changes in mechanical unloading. We found reduced collagen 1 gene expression as well as reduced proline incorporation, an indicator for collagen synthesis, in senescent tissues. Interestingly, Hawthorne *et al* reports increased expression of collagen 1 in aged tendons after treatment with a senolytic therapy (Hawthorne et al., 2023), supporting a relationship between senescence and collagen gene expression. Reduced collagen synthesis could signify inadequate repair of microdamage, loss of mechanical strength, and an inability to adapt to changing mechanical demands. While collagen 1 is the primary structural protein in tendon, proteoglycans and their glycosaminoglycan (GAG) side chains also play a key role in ECM structure and remodeling by promoting water retention and assisting with collagen organization (Siadat et al., 2021). We report differential gene expression profiles of small leucine-rich proteoglycans (SLRPs) decorin and fibromodulin between young and senescent tissues. Interestingly, only young tendons increase proteoglycan expression between days 7 and 14, potentially signifying an adaptive mechanism that is lost in senescent tissues. Furthermore, we report decreased sulfate incorporation, a measure of sulfated GAG synthesis, and decreased GAG content in senescent tissues. As previous work has established a SLRPs as fundamental mediators of tendon healing and repair (Dunkman et al., 2014), as well as regulation of collagen structure, senescence-associated proteoglycan loss could signify inferior tissue remodeling outcomes. Published work in 2D cell culture supports our findings, with both collagen and proteoglycan synthesis reportedly decreasing in senescent fibroblasts (Shelton et al., 1999; Takeda et al., 1992). Additionally, a comprehensive study demonstrates that senescent-induced fibroblasts in an *in vitro* 3D tissue formation model exhibit reduced expression of ECM-related genes as well as reduced collagen fiber deposition and delays in tissue formation (Brauer et al., 2022). Together, this data signifies a compromised ability of senescent cells to synthesize and incorporate new ECM.

We originally hypothesized that senescent tendon explants would display substantial increases in tissue breakdown due to the presence of the SASP phenotype. While ECM degradation is an essential part of routine matrix turnover, extensive breakdown resulting from highly inflammatory states can compromise tissue integrity. However, we did not observe any notable differences between young and senescent groups on the protein level. As discussed previously, stress deprivation injury itself initiates a high inflammatory response in young FDLs, shown by increasing MMP activity and inflammatory protein production. The magnitude of the induced-senescent response is on-par with that of a mechanical injury and sustained throughout the culture period. Interestingly, we found increased expression of MMP-3 and MMP-13 but decreased expression of MMP-1 in senescent tissues compared to young tissues. Prior work cites increased expression of all three enzymes with senescence (Coppé et al., 2010). However, Brauer *et al* report increased MMP-1 expression with DNA damage-mediated senescence but decreased MMP-1 expression in DNA damage-independent senescence compared to controls (Brauer et al., 2022), suggesting induction-specific expression of various SASP factors. Elevated expression of MMP-3, mediator of proteoglycan breakdown, and MMP-13, responsible for both proteoglycan and collagen cleavage (Cui et al., 2017), could explain the lack of GAG accumulation later in culture and signify elevated tissue breakdown in senescent tissues. Our generic MMP activity assay is reflective of nine different types of MMPs, and therefore we did not have individual MMP protein activity in this study. Given our observation that senescence effects different MMPs in distinct ways, future work will look at the production of individual MMPs, as well as tissue inhibitors of MMPs (TIMPs), at the protein level.

Looking at all the data together, we see that compared to young tendons under identical mechanical stimuli, senescent tendons exhibit reduced matrix synthesis with sustained MMP activity, thus shifting the remodeling balance towards degradation over production. A similar conclusion was recently made in a tendon injury model; tendons from a senescent-prone mouse model responded to a collagenase-induced acute injury with differential gene expression profiles from senescent-resistant controls, exhibiting reduced matrix synthesis and increased degradation (Ueda et al., 2019). Interestingly, changes in ECM remodeling were similar between the naturally aged and senescent tendons, suggesting that senescence likely contributes to divergent remodeling outcomes in aged tissues. In some cases, the remodeling response appears graded, with more severe changes in the senescent groups than in aged tissues, possibly due to the fewer number of senescence cells in naturally aged than induced senescent tissues. In other cases, the aged tissue response is no different from senescent tissues, suggesting that the presence of a small number of senescent cells is enough to compromise bulk tissue responses. Ongoing work will assess the impact of altered ECM remodeling on tissue mechanical function, which ultimately affects the ability of the tissue to perform properly *in vivo* and better indicates injury risk.

To our knowledge, this is one of the first studies to induce *in vitro* cellular senescence within the native ECM. This novel senescent explant model allows us to directly assess the contributions of senescent cell populations to altered extracellular matrix remodeling. Importantly, we are able to directly compare the response of senescent tenocytes in 2D cell culture with those remaining in their native microenvironment enabling a more physiologically relevant understanding of the functional consequences of senescence in tendon health. We demonstrate the utility of this model in exploring altered processes of ECM remodeling by investigating how the ECM turnover is disrupted in both naturally aged and senescent-induced tissues following a mechanical unloading injury. We observe that both senescent and aged tendons have a reduced capacity to respond and adapt to altered mechanical environments, likely contributing to the increased prevalence of tendon injuries and poor healing outcomes observed in aging populations. From a clinical perspective, this suggests that senolytic therapies that selectively eliminate senescent cell populations may be a promising strategy for preventing age-related tendon degeneration (Chaib et al., 2022). Our *in vitro* explant model of cellular senescence will give us the opportunity to screen potential treatment strategies and assess the therapeutic ability of senescence-targeting drugs to rescue homeostatic ECM remodeling.

## 5 Acknowledgements

This study was supported by NIH/NIA K99/R00-AG063896 and NSF GFRP (Stowe). Research reported in this publication was supported by the Boston University Micro and Nano Imaging Facility and the Office of the Director, National Institutes of Health of the National Institutes of Health under award Number S10OD024993. The content is solely the responsibility of the authors and does not necessarily represent the official views of the National Institute of Health.

## 8 Authors’ Contributions

EJ Stowe has contributed to all aspects of this study, including research design, data acquisition, interpretation/analysis of data, and drafting/revision of manuscript. MR Keller has contributed to data acquisition, interpretation/analysis of data, and drafting/revision of manuscript. BK Connizzo has contributed significantly to research design, interpretation/analysis of data, and drafting/revision of the manuscript. All authors have read and approved the final submitted manuscript.

