## Supporting_Information for "Cellular Senescence Impairs Tendon Extracellular Matrix Remodeling in Response to Mechanical Unloading"

**Supporting Table S1.** Murine gene names and primer forward/reverse sequences used for quantitative gene expression analysis.

| Target | Gene Name | Forward (5'-> 3') | Reverse (5'-> 3') |
|---|---|---|---|
| Beta Actin | *Actb* [1] | GGCTGTATTCCCCTCCATCG | CCAGTTGGTAACAATGCCATGT |
| p16ink4a | *Cdkn2a* [2] | CGGTCGTACCCCGATTCAG | GCACCGTAGTTGAGCAGAAGAG |
| p53 | *Tp53* [2] | GTACCACCATCCACTACAACTACAT | CAGGACAGGCACAAACACG |
| p21 | *Cdkn1a*[3] | CCTGGTGATGTCCGACCTG | CCATGAGCGCATCGCAATC |
| Lamin B1 | *Lmb1* [4] | GGGAAGTTTATTCGCTTGAAGA | ATCTCCCAGCCTCCCATT |
| Caspase-3 | *Casp3* [5] | ATGGGAGCAAGTCAGTGGAC | CGTACCAGAGCGAGATGACA |
| IL-6 | *Il6* [2] | TAGCTACCTGGAGTACATGAAGAACA | TGGTCCTTAGCCACTCCTTCTG |
| MMP-1 | *Mmp1* [6] | TCAACCAGGCCAAGGTATTG | ATGAGCAGCCACGAGAAATAG |
| MMP-3 | *Mmp3* [5] | ACATGGAGACTTTGTCCCTTTTG | TTGGCTGAGTGGTAGAGTCCC |
| MMP-13 | *Mmp13* [5] | TCAGTCTCTTCACCTCTTTTGGGAATCC | TCAGTTTCTTTATGGTCCAGGCGATG |
| Collagen 1 | *Col1a1* [2] | GACATGTTCAGCTTTGTGGACCTC | GGGACCCTTAGGCCATTGTGTA |
| Decorin | *Dcn* [2] | CTATGTGCCCCTACCGATGC | CAGAACACTGCACCACTCGAAG |
| Fibromodulin | *Fmod* [2] | CTCCAACCCAAGGAGACCAG | GGATCCACCAGTGAGAGTCTTC |

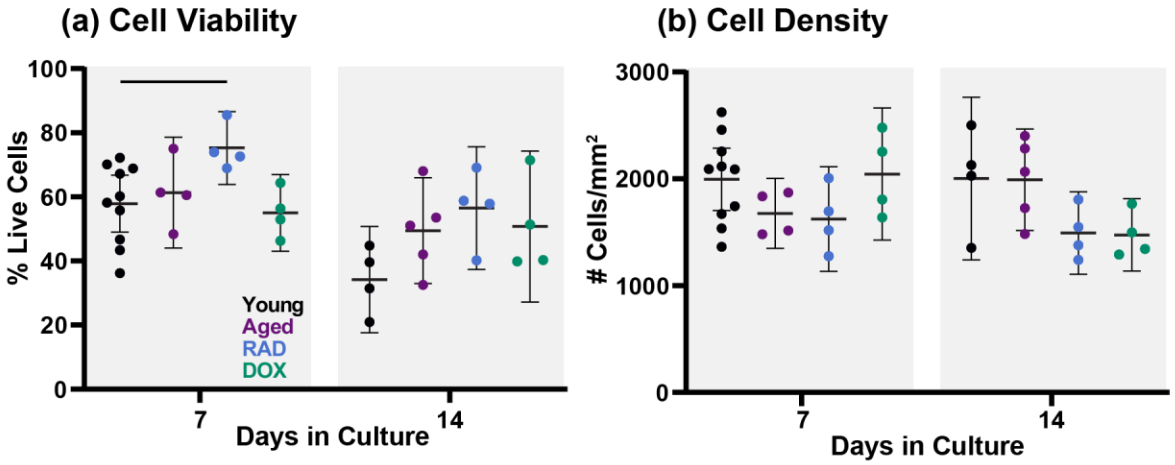


**Supporting Figure S1:** (a) Explant cell viability and (b) cell density quantified from confocal live/dead z-stack images. Statistics show 1-way ANOVAs at each timepoint. Significance set at $p<0.05$ (solid lines) compared to young group.

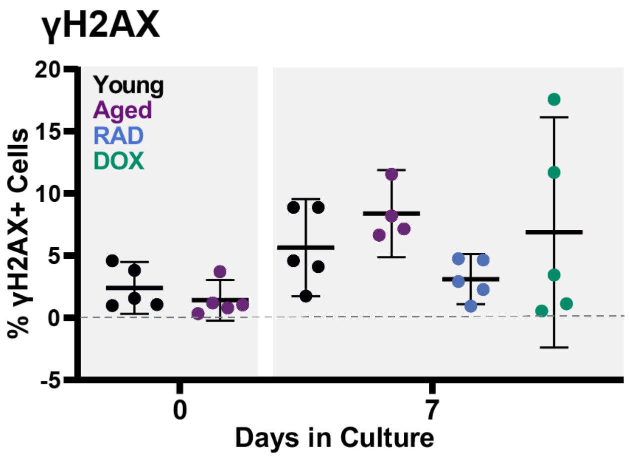


**Supporting Figure S2:** Quantification of γH2AX immunostaining in tendon explants at day 0 and day 7. No statistical differences were found between groups

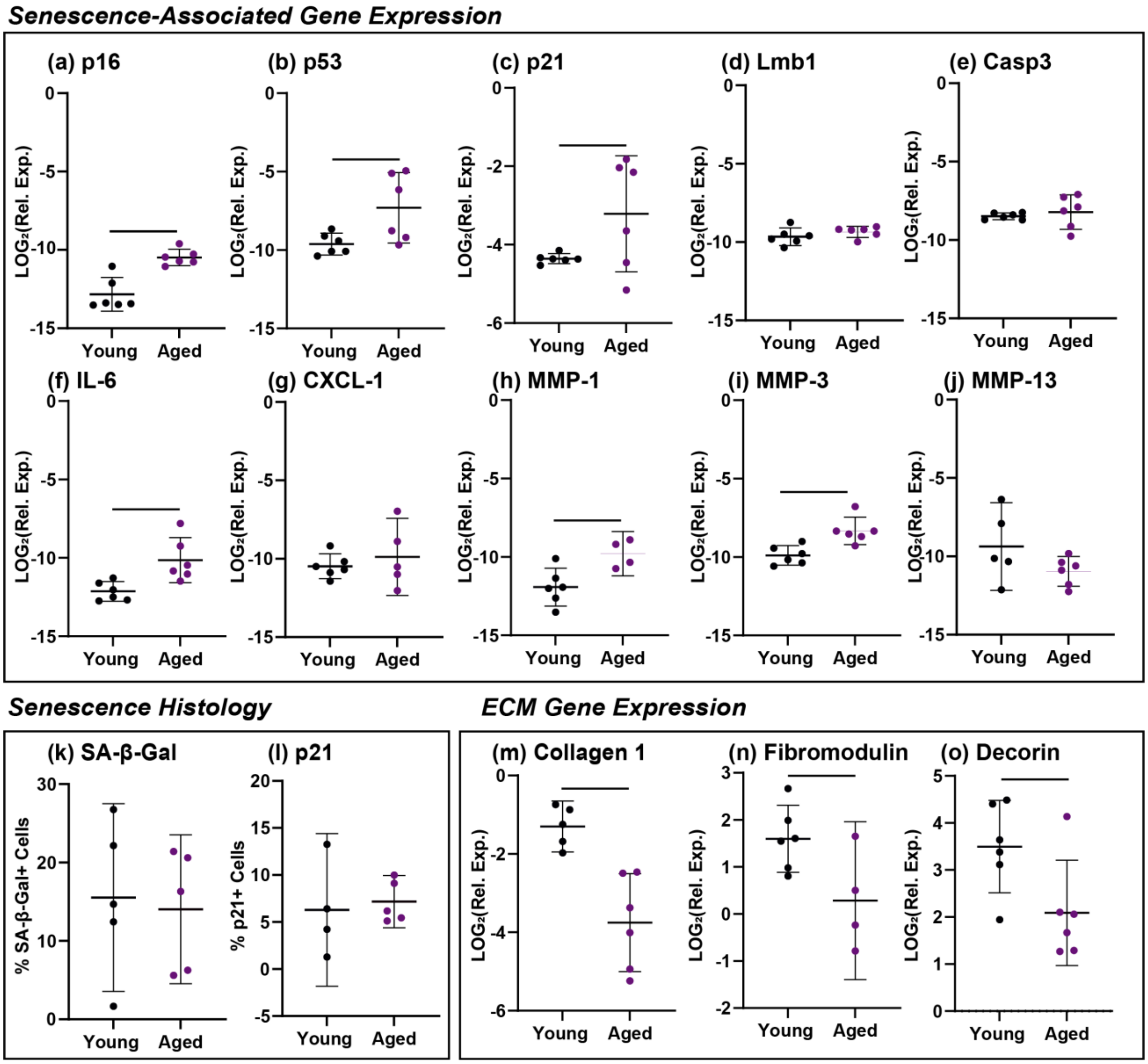


**Supporting Figure S3:** Baseline expression of senescence genes (a) p16, (b) p53, (c) p21, (d) Lmb1, (e) Casp3, (f) IL-6, (g) CXCL-1, (h) MMP-1, (i) MMP-3 and (j) MMP-13. Baseline senescence histology quantifications of (k) SA-β-Gal and (l) p21. Baseline expression of ECM genes (m) collagen 1, (n) fibromodulin, and (o) decorin. Significance set at $p<0.05$ (solid lines) comparing young and aged freshly harvested tissues.
